# DeepPGDB: A Novel Paradigm for AI-Guided Interactive Plant Genomic Database

**DOI:** 10.1101/2025.06.01.657209

**Authors:** Fangping Li, Jiaxuan Chen, Wei Luo, Jieying Liu, Guodong Chen, Binyu Shuai, Zhuangwei Hou, Zhenpeng Gan, Hongyuan Zhao, Penglin Zhan, Changwei Bi, Zefu Wang, Haifei Hu, Shaokui Wang

## Abstract

DeepPGDB (https://www.deeppgdb.chat) is the first AI-driven plant genomics database designed to lower technical barriers in multi-omics research by enabling natural language-based data access and analysis. Integrating over 50 high-quality plant genomes, DeepPGDB combines fine-tuned large language models (LLMs) with prompt engineering to interpret user queries, generate standardized bioinformatics commands, and retrieve or visualize genomic data. Key functionalities include sequence retrieval, BLAST alignment, gene localization, expression profiling, and population genetics analysis, all presented via an intuitive conversational interface. A summarization module further enhances biological reasoning, inferring insights such as haplotype differentiation or protein properties. Benchmarking revealed that Deepseek-r1:14b optimized for short pre-prompts delivers high accuracy and speed. By bridging computational and biological expertise, DeepPGDB democratizes genomic research, fostering interdisciplinary collaboration and accelerating discoveries in agriculture and biotechnology.

---

Dear Editor,

Over the past decade, significant progress has been made in generating omics data, with genomics as a prime example. In plant sciences, high-quality chromosome-level genomes have been published for over 1,000 species (Liu et al., 2024). For model plants such as rice and arabidopsis, advancements have been extended to the population-level genome assemblies, pushing the field into the era of pangenomics. Despite the rapid accumulation of genomic data, researchers with predominantly biological backgrounds often face challenges in mining these datasets and utilizing them as references for multi-omics analyses (Li et al., 2020). These challenges largely stem from entry barriers, including the need for advanced bioinformatics expertise, familiarity with complex command-line tools, programming languages, and data analysis pipelines, as well as intricate user interfaces that require substantial computational skills.

The rapid development of generative large language models, such as ChatGPT and DeepSeek, has recently provided substantial assistance in data processing for researchers (DeepSeek-AI et al., 2025). Furthermore, the emergence of AI-driven agents, such as AutoGPT, has spurred the consideration of placing these models at the core of genomic databases (Yang et al., 2023). In biology, agents have been initially applied to large-scale cancer functional proteomics analysis (Liu et al., 2025). The application and design of artificial intelligence (AI) endow unnatural genomes with desired functions from natural ones (Wang et al., 2025). This advancement has spurred the consideration of placing these models at the core of genomic databases. It is possible to interactively access the rich, embedded knowledge within genome databases through intuitive natural language queries. Guided by this conceptual framework rooted in large language models, our study integrates model fine-tuning with prompt engineering to construct the first AI-powered plant genomics database, DeepPGDB (https://www.deeppgdb.chat). This novel paradigm for AI-guided interactive genomic databases is specifically designed to lower technical barriers, enabling seamless analysis of complex omics data based on high quality genomes. By allowing users from different academic backgrounds to easily access, analyze, and visualize data through natural language queries, DeepPGDB addresses a critical need in the field, paving the way for more productive and inclusive plant genomics research.

Compared to traditional biological data, genomic and multi-omics data are characterized by their large volume and complex formats (Wörheide et al., 2021). Researchers in bioinformatics often rely on specialized tools developed for specific data formats to efficiently extract biologically meaningful information. When adopting AI models as the core scheduler to interactively access data, standardizing the AI model’s output to generate task-specific commands for backend data retrieval and frontend visualization can be an effective approach. In DeepPGDB, we achieve seamless integration of bioinformatics tools by leveraging model integration combined with fine-tuning, prompt engineering, and retrieval-augmented generation (RAG). This enables an AI scheduler to accurately identify user intent and translate it into standardized tool invocation instructions. Task determination in DeepPGDB is approached from two perspectives: task type and data type. First, the AI model interprets user semantics to classify the task as either a tool invocation task or a textual knowledge query task, subsequently selecting the corresponding model to execute the task. For tool invocation tasks, the relevant tool invocation model determines the data type required based on user intent—such as omics data (e.g., expression data, genomic data, or gene function). The AI then generates standardized tool invocation instructions based on a fine-tuned chain-of-thought process and the initial prompt. These instructions are executed on the backend to invoke appropriate tools for comprehensive data retrieval and visualization. The processed results are returned to the frontend for user output (Figure 1A). In contrast, for textual knowledge query tasks (primarily involving stored and related genomic information), the corresponding textual query model employs RAG to retrieve accurate information from relevant documents and provide responses (Figure 1A).

**Figure 1.**
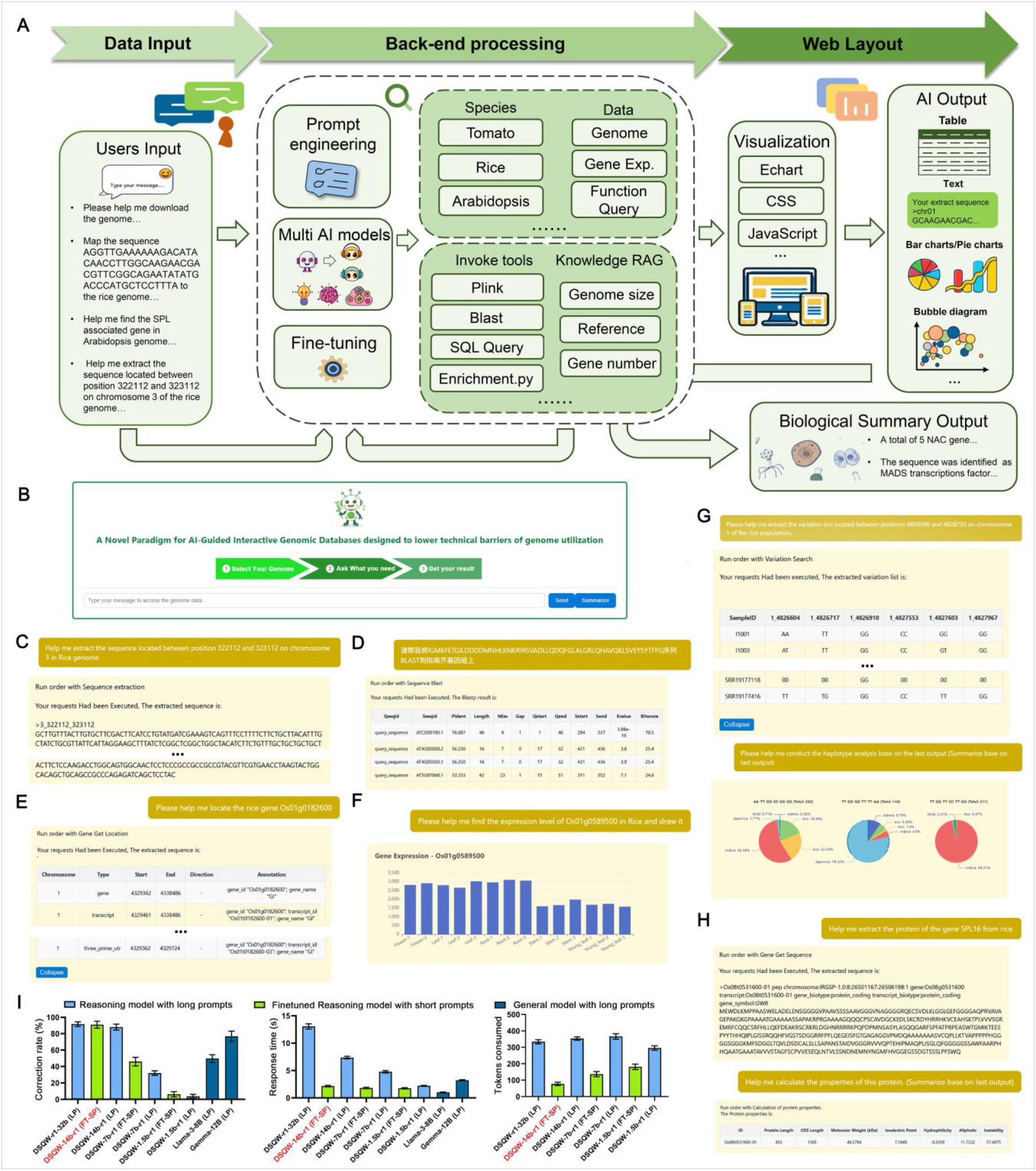
Summary and functions of the DeepPGDB. A: The basic working flow of DeepPGDB; B: The interface of DeepPGDB; C-F: The display example of natural language query and response in basic function of DeepPGDB based on rice and Arabidopsis; C: Sequence extraction for specific intervals from rice genome; D: Protein sequence blast to Arabidopsis with Chinese query; E: Location information of specific genes in rice; F: Visualization of gene expression among various organs in rice; G-H: The display example of advanced summary function of PGDB; G: Variation loci extraction in loci of *GW5L* (Chr01:4826590-4828733) among rice population and the further haplotype analysis based on the output and summarization modula; H: Protein sequence extraction and its basic properties summarize based on the output; Rice *OsSPL16* is taken as an example ; I: The correction rate, response time and tokens consumed of various AI model in DeepPGDB; (DSQW: Deepseek-Qwen; FT: Finetune; SP: Short Pre-Prompt; LP: Long Pre-Prompt).

Users interact with DeepPGDB by submitting queries through a dialog box, with responses processed by the underlying artificial intelligence model and returned in a conversational format (Figure 1A-B). DeepPGDB integrates over 20 publicly available, high-quality plant genomes, enabling users to access and analyze genomic data via natural language input across multiple languages. For example, users can retrieve sequences from specific genomic regions in rice or arabidopsis by submitting queries (Figure 1C; Supplementary Figure 1). Sequence alignment supported by BLAST is a cornerstone for genomic data analysis, which is fully integrated into DeepPGDB. Users can initiate alignment tasks in DeepPGDB by entering nucleotide or protein sequences along with the target species in the dialog interface (Figure 1D; Supplementary Figure 2). The AI model automatically detects the sequence type (nucleotide or protein) and generated standardized backend commands accordingly. Once the commands meet executable standard, the server executes them and returns the results to the frontend in a conversational format.

Genomic location queries and gene list retrieval based on functional categories are crucial for advancing genomic and functional research. In DeepPGDB, we have integrated structural and functional annotation of plant genomes, enabling users to submit queries in natural language that are processed by the AI scheduler. The scheduler interprets user intent and generates standardized Bash or SQL commands, which are executed on the backend to retrieve relevant data. Results are returned and displayed in a structured tabular format on the frontend. For example, when a user queries the chromosomal location of a specific rice gene, the AI automatically extracts the rice gene identifier, formulates a corresponding standardized backend command, executes it, and displays the results on the frontend (Figure 1E). Similarly, for gene family queries, the AI identifies relevant keywords and searches the backend and returns a comprehensive gene list (e.g., the GRF transcription factor family in rice; Supplementary Figure 3). To accommodate species with multiple annotation versions, DeepPGDB employs a multi-annotation correspondence table in the backend. In the case of rice, this includes mappings across various identifiers such as RAP (e.g., *Os02g0661100*), MSU (e.g., *LOC_Os02g44230*), and gene names (e.g., *OsTPP1*). Upon receiving a query, the AI matches the terms provide by users to these identifiers. If a unique query is found, the system returns the results directly. However, if multiple entries are identified, the DeepPGDB presents all relevant candidate genes along with their annotation, prompting users to refine the query using a specific gene ID (Supplementary Figure 4).

In addition to basic queries and tabular outputs, DeepPGDB integrates interactive statistical visualization tools powered by ECharts, enabling users to intuitively explore data through dynamic charts. Gene expression profiles curated for the included species have been integrated into DeepPGDB, allowing users to visualize differential expression patterns between groups within a given dataset using natural language instructions (Figure 1F). Similar visualization and analysis workflows are available for enrichment analyses based on species-specific gene lists (Supplementary Figure 5). Additionally, population genetics, a critical area in plant research, is also supported by DeepPGDB through the integration of population genomic variation data from multiple species. Upon receiving user’s requests, the AI interprets the query, invokes the PLINK tool (Purcell et al., 2007) retrieves results from the preloaded population datasets. These outputs are then presented in a structured tabular format on the frontend (Figure 1G).

DeepPGDB’s interactive and user-friendly framework for omics data retrieval and visualization meets the fundamental expectations of modern biological databases. However, contemporary large language models offer capabilities that extend beyond basic data querying, enabling more advanced reasoning and synthesis. Inspired by knowledge bases like Tencent IMA.Copilot enable models to extract information from text and generate logical summaries and inferences; we introduced a summarization module into DeepPGDB. This module serve as a reasoning layer that infers potential biological significance based on user requests and the results of prior interactions. Its inference capabilities are supported by a foundational knowledge framework, scientific reasoning logic obtained through fine-tuning, and prompt engineering, and structured prompt engineering. For example, the module can summarize the haplotypes of the *GW5L* (Tian et al., 2019) genomics intervals across rice subspecies, revealing subspecies-specific haplotype differentiation (Figure 1G). In another use case, the summarization module can extract the sequence of the rice gene *OsSPL16* (Wang et al., 2012) and calculate its protein properties (Figure 1H), illustrating its capacity to perform multi-step biological reasoning and analysis.

The size of parameters for generative models is closely related to their reasoning capabilities. Under specific hardware constraints, models with appropriately sized parameters and task-specific reasoning architectures can effectively balance generation efficiency and task performance. To optimize DeepPGDB performance in practical applications, we benchmarked several candidate core models under identical quantization settings and hardware conditions (a single V100-SXM2 GPU with 32 GB VRAM combined with a dual-processor system utilizing AMD EPYC 7642 CPUs) (Figure 1I). We evaluated each model in terms of output accuracy and reasoning time. The comparison showed that the 14-billion-parameter reasoning model (Deepseek-r1:14b) achieved optimal performance, achieving approximately 90% accuracy under the Long Pre-prompt (LP; length about 2100 tokens Supplementary File 1) across various tasks, while maintaining superior response speed compared to larger reasoning models (e.g., 32b). In contrast, a similarly scaled general-purpose model (Gemma:12b) exhibited significantly faster instruction generation speed but exhibited notably lower accuracy in instruction generation due to its absence of reasoning capabilities. To further enhance performance, we fine-tuned the reasoning model to support fast reasoning under the Short Pre-prompt (SP; length about 400 tokens Supplementary File 1) with reduced token consumption while maintaining the high accuracy observed under LP. This optimization significantly improved responsiveness, making it more suitable for deployment in DeepPGDB. We also tested smaller reasoning models (7b/1.5b) which showed modest accuracy gains following fine-tuning under the same corpus. However, these models failed to meet performance requirements, with frequent hallucinations and diminished reasoning accuracy, despite lower token usage compared to other larger parameter models (14b) (Figure 1I).

The introduction of DeepPGDB marks a transformative step in the field of plant genomics, fostering interdisciplinary collaboration and bridging gaps between computational biology, genomics, and agricultural sciences. By integrating high-quality genomic and multi-omics data with advanced AI-driven scheduling, standardized command generation, and interactive visualization tools, DeepPGDB enables researchers from diverse backgrounds to efficiently access and interpret complex biological data. This nuanced approach not only supports precise gene localization, functional annotation, and population genetics studies but also enhances the extraction of biologically relevant insights through its summarization module. Although still in its early stages, our focus is on the systematic collection of high-quality plant genomic data to ensure the database remains comprehensive and current. We also plan strategic updates to our prompt engineering framework—such as fine-tuning existing models and exploring larger-scale integrations—to broaden task capabilities and enhance the extraction of biologically relevant insights. Ultimately, by facilitating a deeper understanding of genetic data, DeepPGDB is poised to drive significant breakthroughs in agricultural science, conservation, and biotechnology, thereby propelling the field toward a new era of data-driven discovery.

