## Supplemental Figures for "DeepPGDB: A Novel Paradigm for AI-Guided Interactive Plant Genomic Database"

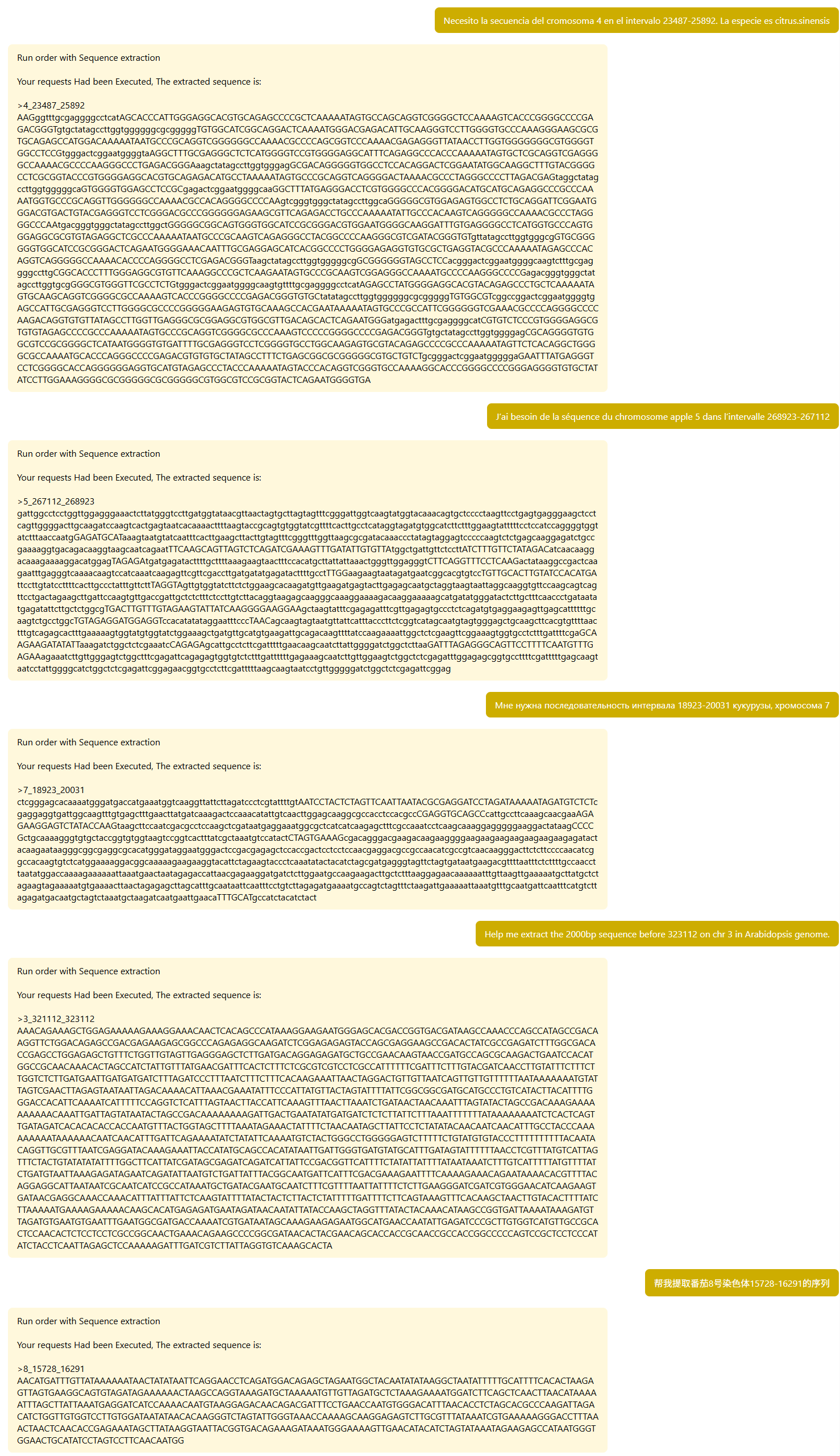

**Supplemental Figure 1. Sequence extraction in multilingual (French, Spanish, Russian, English, Chinese) and multispecies (orange, apple, corn, Arabidopsis, tomato) cases from DeepPGDB**

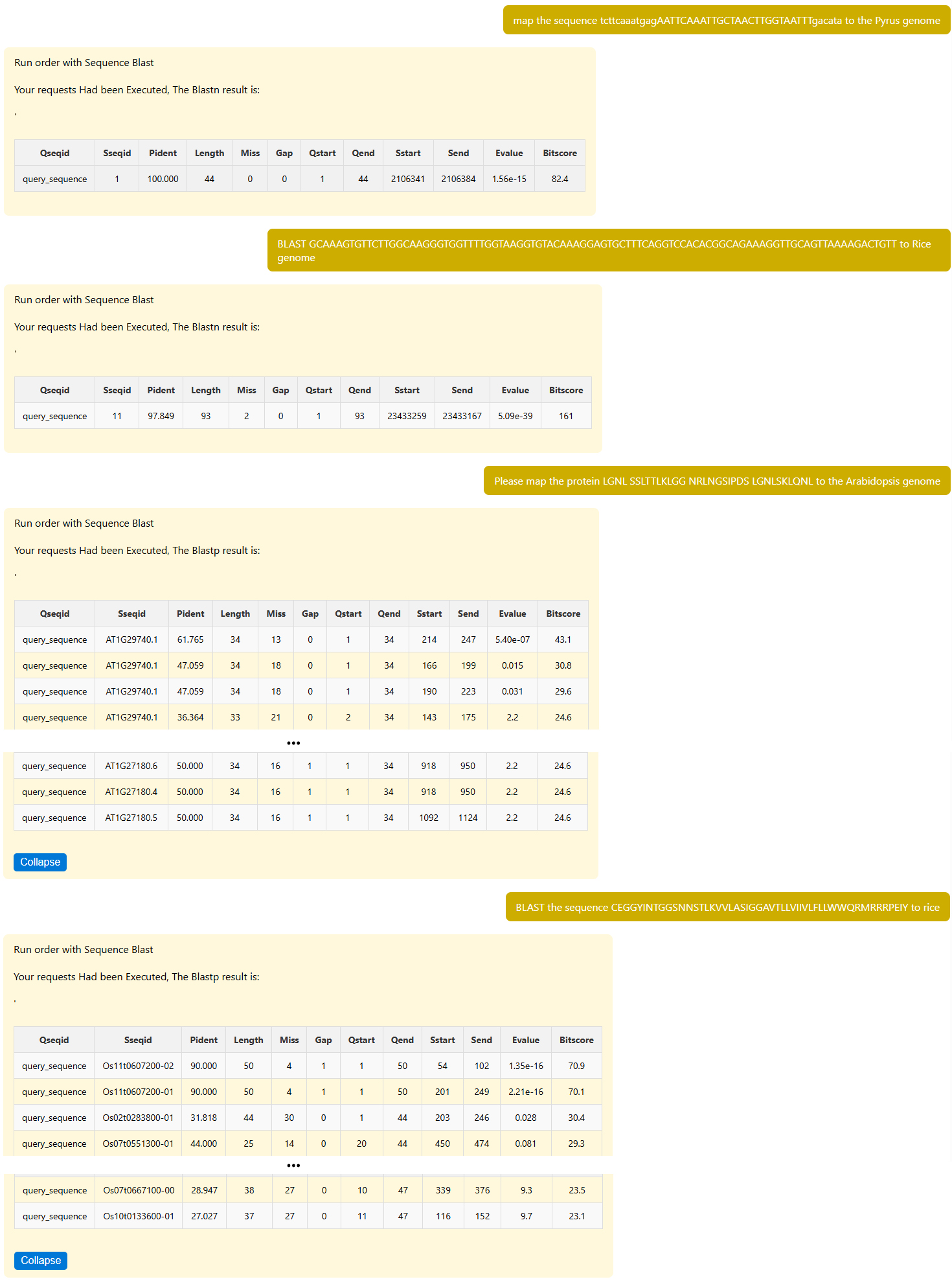

**Supplemental Figure 2. Nucleotide BLAST (Blastn) and protein BLAST (Blastp) applied in multispecies (pear, Arabidopsis, rice) from DeepPGDB**

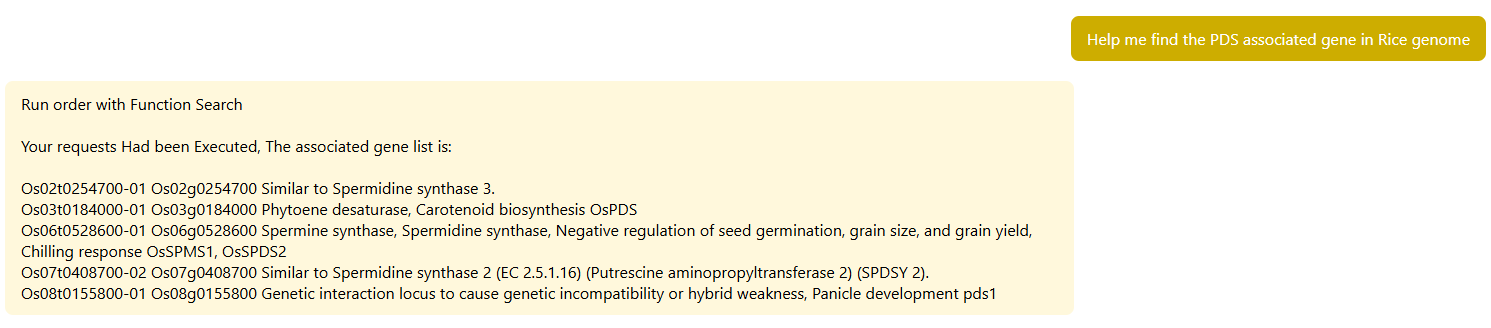

**Supplemental Figure 3. Gene information search of PDSs in rice genome** **from DeepPGDB**

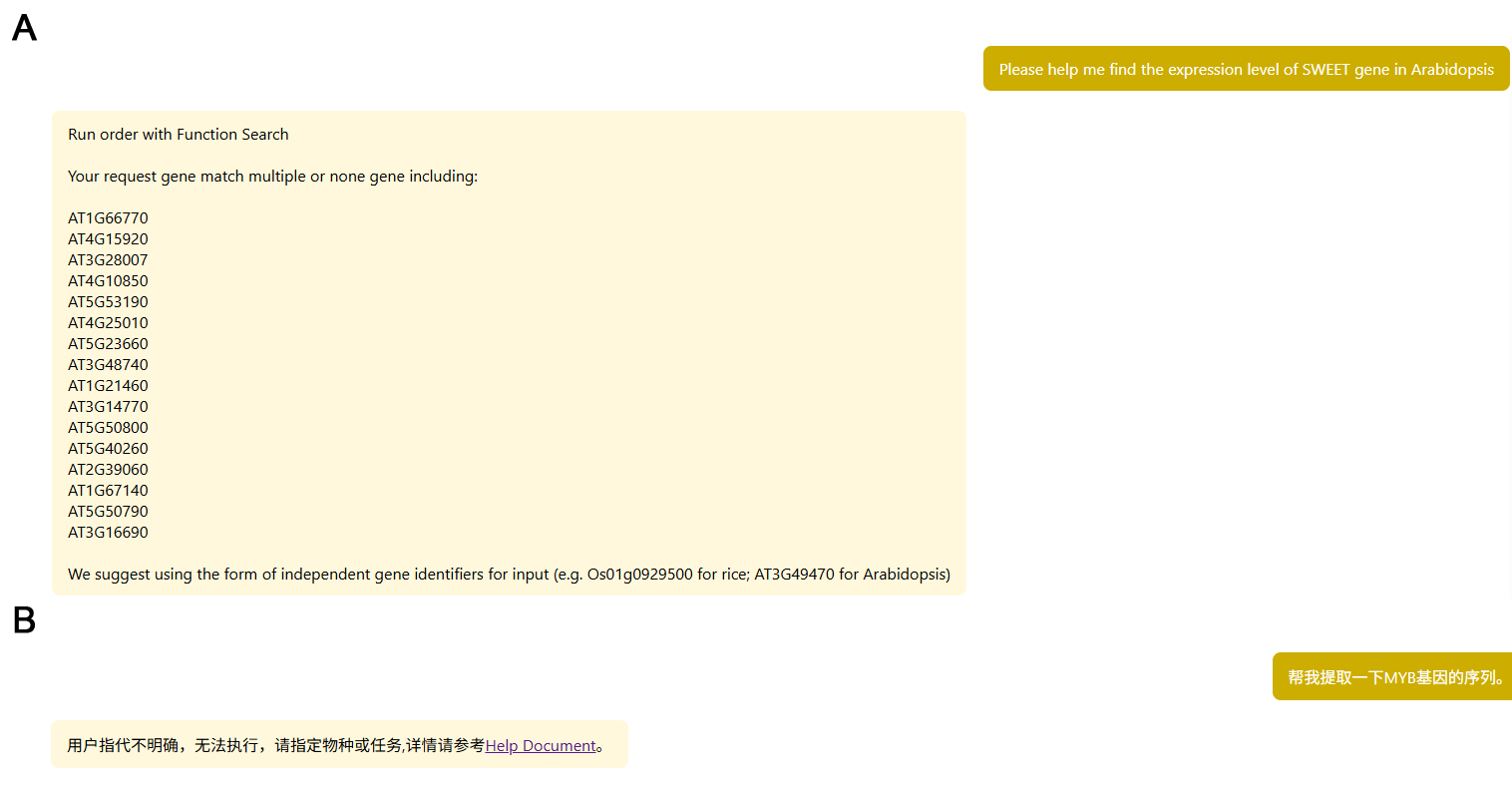

**Supplemental Figure 4. Two solutions for low-confidence user input from DeepPGDB. A: Output points to multiple results; B: Incorrect species or tasks (in Chinese)**

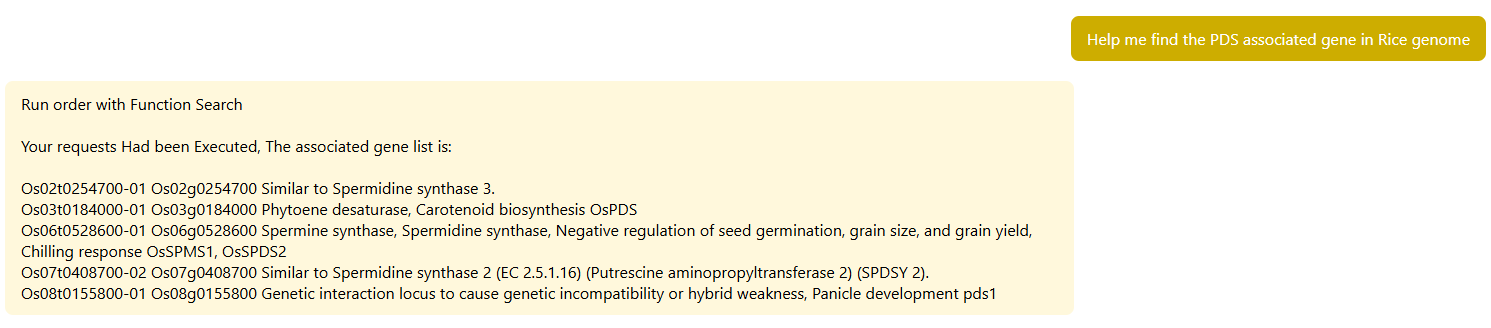

**Supplemental Figure 5: GO enrichment analysis and visualization in Arabidopsis from DeepPGDB**
