## Supplemental Files for "DeepPGDB: A Novel Paradigm for AI-Guided Interactive Plant Genomic Database"

**Supplemented File 1: Method**

***Web-server deployment***

The front-end and back-end interaction of deepPGDB is accomplished by the Python-based Flask web framework (https://flask.palletsprojects.com/zh-cn/stable/). The back-end response service is handled by Nginx (https://nginx.org/en/). The domain name and IP obtain SSL certificates from the Let's Encrypt platform through Cerbot (https://certbot.eff.org/).

***Finetune of the model***

We fine-tuned three backbone models (DeepSeek-R1-Distill-Qwen-7B, DeepSeek-R1-Distill-Qwen-1.5B and DeepSeek-R1-Distill-Qwen-14B) (DeepSeek-AI et al.,2025) on the LLaMA Factory platform (Zheng et al.,2024) using the Quantized Low-Rank Adaptation (QLoRA) (Dettmers, Pagnoni, Holtzman, & Zettlemoyer,2023) framework with 4-bit quantization. Training was performed on NVIDIA A100 PCIE 40G GPU using a curated training set (https://github.com/lifangpings/DeepPGDB/data/primary.training.json) with more than 300 question–answer pairs with AdamW (https://optimization.cbe.cornell.edu/index.php?title=AdamW) optimizer in parameters of learning rates=5.0e-6, batch sizes=6, number of epochs=3).

***Identification and processing of potential ambiguities***

We enhanced the core model's understanding of its own capabilities through prompt engineering, and achieved this by adding noise to the fine-tuning training set (consistent with the model's judgment when encountering noisy inputs), enabling it to guide users back to tasks within their capabilities (such as guiding users to return to tasks within their range, for example, by providing input examples for explanation) when faced with low confident query

***The Measurement of model performance***

All the models were and tested on the computing platform of (a single V100-SXM2-32G card combined with AMD EPYC 7642). All the models have undergone Q4_K_M quantization in the llama.cpp (https://github.com/ggml-org/llama.cpp). In the benchmark tests, the models uniformly performed various tasks including sequence query, sequence alignment, expression quantity query, gene location information, query visualization, etc. The benchmark tests consisted of n groups (n > 10), and each group conducted five to six conversations.

***Data available***

The details of question example and latest database data scope is available in the description document of the database (<https://deeppgdb.chat/document.html>) The fine-tuned models generated in DeepPGDB, along with the adapted interface scripts, front-end web files, back-end working scripts and Pre Prompt including long Pre Prompt and Short Prompt, were respectively uploaded to Modelscope (https://www.modelscope.cn/models/LEECHXP/DeepPGDB) and GitHub (https://github.com/lipingfangs/DeepPGDB).
